# Influence of short-term temperature drops on sex-determination in sea turtles

**DOI:** 10.1101/2021.03.29.437634

**Authors:** Ellen Porter, David T. Booth, Colin J. Limpus

## Abstract

All sea turtles exhibit temperature-dependent sex-determination, where warmer temperatures produce mostly females and cooler temperatures produce mostly males. As global temperatures continue to rise, sea turtle sex-ratios have become increasingly female-biased, threatening the long-term viability of many populations. Nest temperatures are dependent on sand temperature, and heavy rainfall events reduce sand temperatures for a brief period. However, it is unknown whether these short-term temperature drops are large and long enough to produce male hatchlings. To discover if short-term temperature drops within the sex-determining period can lead to male hatchling production, we exposed green and loggerhead turtle eggs to short-term temperature drops conducted in constant temperature rooms. We dropped incubation temperature at four different times during the sex-determining period for a duration of either 3 or 7 days to mimic short-term drops in temperature caused by heavy rainfall in nature. Some male hatchlings were produced when exposed to temperature drops for as little as 3 days, but the majority of male production occurred when eggs were exposed to 7 days of lowered temperature. More male hatchlings were produced when the temperature drop occurred during the middle of the sex-determining period in green turtles, and the beginning and end of the sex-determining period in loggerhead turtles. Inter-clutch variation was evident in the proportion of male hatchlings produced, indicating that maternal and or genetic factors influence male hatchling production. Our findings have management implications for the long-term preservation of sea turtles on beaches that exhibit strongly female-biased hatchling sex-ratios.

## INTRODUCTION

Sea turtles spend their entire life in the ocean, except for when females return to the beach to nest (Miller, 1997; Spotila, 2004). Sea turtle embryonic development is dependent on nest temperature which influences incubation time, hatching success, hatchling size, hatchling sex, and hatchling locomotion performance (reviewed in Ackerman, 1997; Booth 2017). The sensitivity of sea turtle embryos to temperature make these animals particularly susceptible to global warming, with numerous studies suggesting global warming could result in severe declines in sea turtle populations due to increases in incubation temperature (Mrosovsky and Provancha, 1992; Ozdemir et al., 2011; Fuentes et al. 2012; Katselidis et al., 2012; Patino-Martinez et al., 2012; Saba et al., 2012; Weber et al., 2012; Fisher et al., 2014; Wood et al., 2014; Jensen et al., 2018; Booth et al., 2020; Reboul et al., 2021).

All sea turtles exhibit temperature-dependent sex-determination (TSD), where sex is determined by temperature during the sex-determining period (SDP) (Yntema and Mrosovsky, 1980; Standora and Spotila, 1985; Wibbels, 2003). The SDP occurs during the middle third of embryonic development when the gonads are differentiating, with cooler temperatures producing more males and warmer temperatures producing more females (Yntema and Mrosovsky, 1982; Miller and Limpus, 1981; Limpus et al., 1985; Standora and Spotila, 1985; Godfrey and Mrosovsky, 2006; Woolgar et al., 2013). The transitional range of temperatures (TRT) is range of temperatures over which sex-ratios shift from 100% male to 100% female (Fig. 1). The pivotal temperature occurs within the TRT, and is the incubation temperature at which a one-to-one sex-ratio is produced (Fig. 1) (Girondot, 1999; Wibbels, 2003). For example, the TRT range and pivotal temperatures for the southern Great Barrier Reef (sGBR) green turtle (*Chelonia mydas*) population are 27.4 – 28.7 °C and 28.1 °C respectively and for the East Australian (EA) population of loggerhead turtles (*Caretta caretta*) they are 25.0 – 32.0 °C and 28.4°C respectively (Fig. 1).

**Figure 1.**
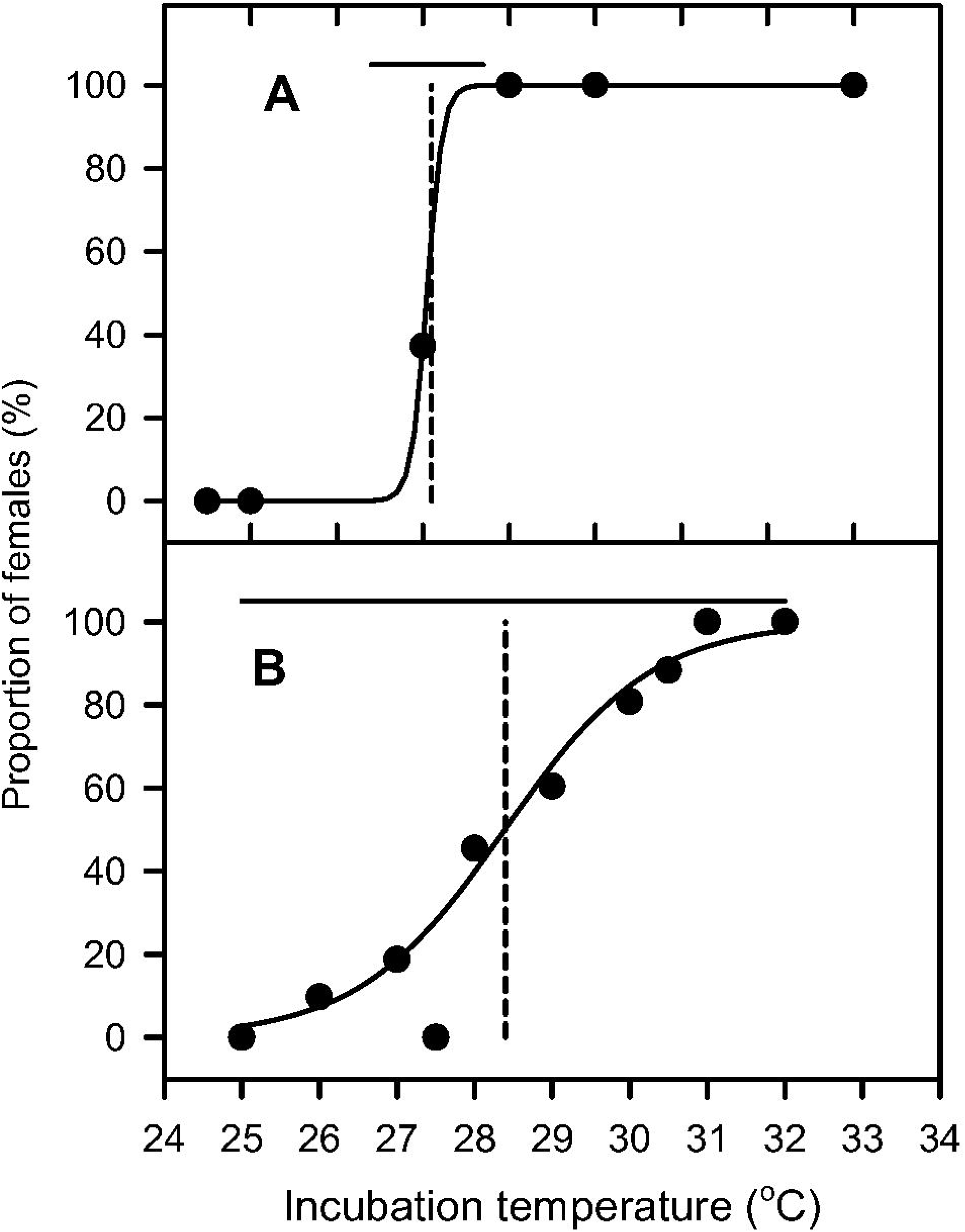
Relationship between incubation temperature and proportion of female hatchlings for (A) the southern Great Barrier Reef (sGBR) nesting population of green turtles, and (B) the East Australian (EA) nesting population of loggerhead turtles. Lines fitted to data with logistic regression of the form: Proportion of females = (e^y^)/(1 + e^y^), where y = b x incubation temperature + c. For sGBR green turtles b = 11.1422 and c = −312.5039. For EA loggerhead turtles b = 1.0568 and c = −30.0023. Solid horizontal line indicates the transitional temperature range, dashed vertical tine indicates the pivotal temperature. Data for sGBR green turtles obtained from Miller and Limpus (1981) and Burgess et al. (2006). Data for EA loggerhead turtles obtained from Limpus et al. (1985).

Hatchlings sea turtles lack sexual dimorphism and heteromorphic sex chromosomes (Wibbels, 2003; Ceriani and Wyneken, 2008), so histological examination of the gonad, which involves the euthanasia of the hatchlings, has been the only reliable method to determine their sex (Yntema and Mrosovsky, 1980; Miller and Limpus, 1981; Wibbels, 2003; Ceriani and Wyneken, 2008). Recently, a method for determining sea turtle hatchling sex using a small blood sample was described (Tezak et al., 2020), but this method is yet to be verified across all sea turtle species. Estimation of sea turtle hatchling sex-ratio using the temperature during the SDP is frequently used in lieu of histology because of a reluctance to euthanise hatchling sea turtles to obtain their gonads (Yntema and Mrosovsky, 1980; Booth and Astill, 2001b; Chu et al., 2008; Wood et al., 2014; Laloë et al., 2016). This method involves fitting mathematical models to empirical sex-ratio data gathered from constant temperature incubation experiments where hatchling sex was determined by histology (Fig. 1). However, natural nest temperatures are never constant, so determining the middle third of embryonic development when sex is determined does not correspond with the middle third of the incubation period. To solve this problem, a mathematical model can be fitted to constant temperature - incubation period data (Fig. 2) and the inverse of this relationship used to predict the embryonic development rate at any temperature. By summing together the developmental increments for each hour of development determined from nest temperature traces, the stage of embryonic development at any point in time of incubation can be calculated (Booth and Freeman, 2006). Once the middle third of embryonic development is determined using this method, the constant temperature equivalent (CTE) (Georges et al., 1994) during the SDP can be calculated. This CTE can then be substituted into the algorithm relating temperature to hatchling sex-ratios to predict the hatchling sex-ratio for any nest where incubation temperature has been continuously monitored (Booth and Freeman, 2006; Chu et al., 2008).

**Figure 2.**
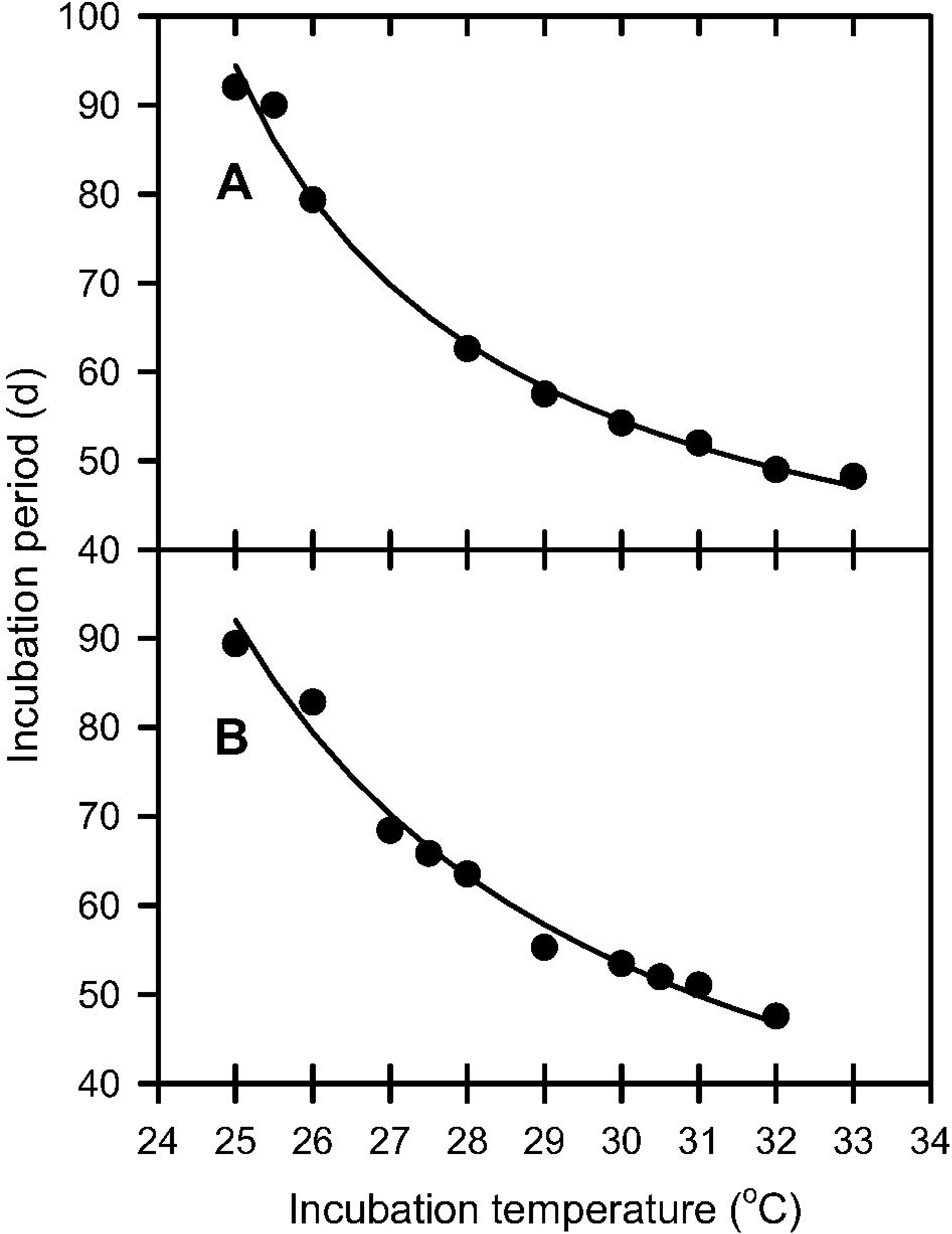
Relationship between constant incubation temperature and incubation period for (A) the southern Great Barrier Reef (sGBR) nesting population of green turtles, and (B) the East Australian (EA) nesting population of loggerhead turtles. Lines fitted to data with inverse regression of the form: Incubation period = c + (a/(incubation temperature + b). For sGBR green turtles, a = 239.7053, b = −21.4777 and c = 26.3928. For EA loggerhead turtles, a =417.2294, b = −19.7319 and c = 12.8043. Data for sGBR green turtles obtained from Bustard and Greenham (1968), Miller and Limpus (1981), Miller 1982, Booth and Astill (2001a), Booth et al. 2004, and Burgess et al. (2006). Data for EA loggerhead turtles obtained from Limpus et al. (1985).

Increased nest temperatures caused by global warming result in strongly female-biased sex-ratios which may threaten the long-term persistence of many sea turtle populations (Reed, 1980; Mrosovsky and Provancha, 1992; Godfrey et al., 1996; Broderick et al., 2000; Booth and Astill, 2001b; Patino-Martinez et al., 2012; Wood et al.,2014; Laloë et al., 2016; Jensen et al., 2018). Cooling techniques such as shading with natural vegetation, relocating eggs into shaded hatcheries or irrigating nests are management actions used at sea turtle nesting beaches to reduce nest temperatures in order to increase male hatchling production (e.g. Patino-Martinez et al., 2012; Wood et al., 2014; Hill et al., 2015; Jourdan and Fuentes, 2015; Esteban et al., 2018).

Nest temperature is determined by sand temperature (heated via solar radiation) and metabolic heat production of the developing embryos. Sand temperature is dependent on beach orientation, sand colour and the presence of absence of shade (Broderick et al., 2001; Hays et al., 2001; Ackerman and Lott, 2004; Booth and Freeman, 2006; Wood et al., 2014; Hill et al., 2015). Sand temperature also fluctuates over a daily heating/cooling cycle; however, the magnitude of this cycle decreases as sand depth increases (Georges et al., 1994; Booth and Freeman, 2006; Booth, 2006). Sand temperature is also influenced by latitude. At higher latitudes, where there is strong seasonal heating and cooling of sand throughout the year, nest temperatures are slightly warmer at shallower depths during the sand heating period (beginning of summer), but slightly cooler during the sand cooling period (end of summer) (Godley et al., 2001; Booth and Freeman, 2006). Conversely, nests located closer to the equator may not experience noticeable seasonal fluctuations in sand temperature (Spotila et al., 1987). Nest temperature increases 1-3 °C above sand temperature towards the end of incubation due to metabolic heat produced by the embryos as they grow larger in size (Booth and Astill, 2001b; Broderick et al., 2001; Booth and Freeman, 2006; Booth, 2017; Rivas et al., 2019). However, during the SDP, metabolic heating has little effect on nest temperature because the embryos are still small and produce negligible metabolic heat (Booth and Astill, 2001b; Broderick et al., 2001).

While cooler nest temperatures increase male hatchling production, it is unknown whether short-term drops in nest temperature caused by heavy rainfall can induce male hatchling production from beaches that would otherwise produce only female hatchlings. Heavy rainfall events cause sand and nest temperatures to drop between 2°C and 4°C for several days post-rain (Reed, 1980; Booth and Astill, 2001b; Booth and Freeman, 2006; Wood et al., 2014; Sim et al., 2015; Rivas et al., 2018; Staines et al., 2019, 2020; Laloë et al., 2020; Reboul et al., 2021). The duration of the temperature drop after heavy rainfall is highly variable, from just two to three days if warmer weather returns immediately after rain, to more than a week if rainy weather and/or heavy cloud persists for several days after the initial rainfall event. In the laboratory, short-term drops in incubation temperature (5-10 days) in loggerhead turtle eggs during the SDP were able to induce some male hatchling at otherwise all female incubation temperatures (Woolgar et al., 2013). Our study aims to extend this study to discover if exposure to cool temperatures for as short as three days can induce male hatchling production. We used green and loggerhead turtle eggs in a series of temperature drop experiments that mimic the drops in temperature that are experienced after heavy rainfall events. These experiments determined if the duration of the temperature drop (3 or 7 days), and the timing of the temperature drop within the SDP can result in male hatchling production. If our experiments result in male production, these finding can be used as a guide to sea turtle nesting beach managers who are interested in increasing male hatchling production at their beach, by demonstrating that one-off irrigation of nests that reduce nest temperature by 3 °C can result in male hatchling production.

## MATERIAL AND METHODS

### Study sites, egg collection and transport

Four clutches of the sGBR nesting population of green turtle eggs were collected on 12^th^ January 2020 from Heron Island, Great Barrier Reef (23°26′S, 151°55′E). Eggs were placed into plastic bags (one clutch per bag) for transportation to Heron Island Research Station (10 minute walk), then placed into an insulated container with the lid left open in a 10°C room until departure the next day. Lowering the temperature pauses embryonic development so that movement-induced mortality is reduced during transit (Parmenter, 1980). The container was closed one hour prior to departure to maintain a low temperature during transit. Following a boat trip to Gladstone, eggs were transported by car to the laboratory at The University of Queensland, St Lucia campus. The entire transportation process took 9 hours.

One clutch of the EA nesting population of loggerhead turtle eggs was collected on 6^th^ February 2020 from Mon Repos beach on the east coast of Australia (24°48’S, 152°26’E). The eggs were chilled to ~10°C in a transportable refrigerator (Engel 40L; MT45F-S) overnight before being transported by car to the laboratory at The University of Queensland, St Lucia campus. The entire transportation process took 6 hours.

All experiments were approved by The University of Queensland NEWMA animal ethics committee, certificate number AE56601. Green turtle eggs were collected under permit PTU19-002268 issued by Queensland Government Parks and Wildlife Service to DTB. Loggerhead turtle eggs were collected under a management directive authorised by CJL.

### Experimental design

Sea turtle eggs were incubated in two constant temperature rooms, one set at 30.0°C (a female producing temperature), the other at 26.5°C (a male producing temperature). Most eggs were incubated predominantly at 30.0°C, and eggs temporarily moved to 26.5°C for either 3 or 7 days at different times during the SDP and then moved back to 30.0°C (Table 1). A sample of eggs from each clutch was used in each treatment, but the number of eggs from each clutch varied because the total number of eggs varied between clutches. The duration of the 26.5°C exposure was chosen to mimic a single short heavy rainfall event and a more prolonged rainfall event with associated cloudy weather that occur in nature, respectively. The start and end of each cooling period was staggered across the SDP to determine if male production might be dependent on the timing of temperature shifts within the SDP. The incubation period for green turtle eggs incubated at 30.0°C is 56 days (Fig. 2A), with the SDP occurring between day 19 and 37. For loggerhead turtle eggs, the incubation period at 30.0°C is 54 days (Fig. 2B), with the SDP occurring between day 18 and 36 of incubation.

**Table 1.** The 10 experimental treatments used to incubate green and loggerhead turtle eggs.Treatments 1 and 2 were male and female controls which were incubated continuously at a constant temperature of 26.5°C and 30.0°C, respectively. Eggs in treatments 3–10 started incubating at 30.0°C before being moved to 26.5°C for either 3 or 7 days, then moved back to 30.0°C for the remainder of incubation. Although treatments started incubation at the same time, clutches were shifted to 26.5°C at different times throughout the incubation period, with temperature drops starting between day 19 (the beginning of the sex-determining period) and day 30 (a week before the end of the sex-determining period).

| Treatment | Day exposure to<br>26.5°C began | Drop duration (days) | Day exposure to<br>26.5°C ended |
| --- | --- | --- | --- |
| T1 | Male control (26.5°C throughout incubation) |  |  |
| T2 | Female control (30.0°C throughout incubation) |  |  |
| T3 | 19 | 3 | 22 |
| T4 | 19 | 7 | 26 |
| T5 | 23 | 3 | 26 |
| T6 | 23 | 7 | 30 |
| T7 | 27 | 3 | 30 |
| T8 | 27 | 7 | 34 |
| T9 | 30 | 3 | 33 |
| T10 | 30 | 7 | 37 |

### Egg processing and preparation

On arrival in the laboratory, eggs were rinsed in freshwater to remove adhering sand and a unique identification number showing the clutch of origin and individual number written on the top surface of the egg using a 2B pencil. Eggs from each clutch were split across 10 different containers (between 9 and 12 eggs per container depending on clutch size) representing each of the 10 treatments and were covered with sterilised Mon Repos beach sand mixed with distilled water (6 grams of water per 100g of dry sand) to give a water potential of ~ −100kPa. An iButton (DS19221L, Maxim Integrated, San Jose, California, USA) temperature data logger, set to log temperature every hour, was placed amongst the eggs and plastic cling-wrap was placed over the top of the containers to reduce the rate of water lost from the sand. The containers were placed on racks in the 30.0°C constant temperature room, or in the case of treatment T1, in the 26.5°C constant temperature room. To prevent the effect of location within the constant temperature rooms being misinterpreted as clutch effects (Wibbels, 2003), the containers were randomly assigned different locations within the room and moved from one shelf to another every week.

Forty-eight hours into incubation, sand covering the top surface of the eggs was brushed aside to check for the formation of a “white-patch” that indicates embryonic development had begun. Eggs not displaying a white-patch were removed. We checked containers weekly to identify and remove dead eggs to prevent any fungus/disease from spreading. During this time, we also added distilled water to the sand via a spray bottle to replace the moisture in the sand that had been absorbed by the eggs or lost through evaporation.

On their prescribed day, containers of eggs in treatments 3 to 10 (Table 1) were moved into the 26.5°C room for their temperature drop duration (3 or 7 days). Once their temperature drop period concluded, containers were moved back into the 30.0°C room. From day 48 of incubation onwards we inspected containers every 6 hours for eggs that had pipped indicating the beginning of hatching. Pipped were removed and placed in isolated containers within the incubator until hatching was completed.

### Hatchling gonad histology and identification of sex

Once hatchlings emerged from their eggs, they were euthanised by an overdose of a gaseous anaesthetic (2-4 ml isoflurane inside a sealed container) for 1 hour. After death, each hatchling was dissected and the left kidney-gonad complex removed and fixed in a 10% buffered formalin solution for 24–48 hours before being stored in a 70% ethanol solution until prepared for histology. For histology, the kidney-gonad complexes were placed into cassettes and subsequently dehydrated using a Leica^©^ ASP300S tissue processor before being embedded in paraffin wax blocks. The blocks were cut into 6 µm sections using a Leica^©^ rotary microtome and mounted onto slides. These slides were stained using the hematoxylin and eosin staining process (Bancroft and Layton, 2012). Cross-sections of gonad and associated ducts were digitally scanned using a Leica^©^ Aperio ScanScope XT brightfield microscope slide scanner. Images were analysed using Aperio ImageScope imaging software.

A hatchling’s sex was determined by the presence of either ovary or testis tissue as described for sea turtle hatchlings (Yntema and Mrosovsky, 1980; Miller and Limpus, 1981; King et al., 2013; Sari and Kaska, 2016). Females had ovaries characterised by a thickened cortical layer, with the medulla consisting of dense mesenchymal tissue (Fig. 3A), as well as the Müllerian duct showing a columnar epithelium and lumen with a long stalk (Fig. 3B). Males had testes having little to no cortex and differentiated seminiferous tubules within in the medulla (Fig. 3C) and a degenerated Müllerian duct with a short stalk (Fig. 3D).

**Figure 3.**
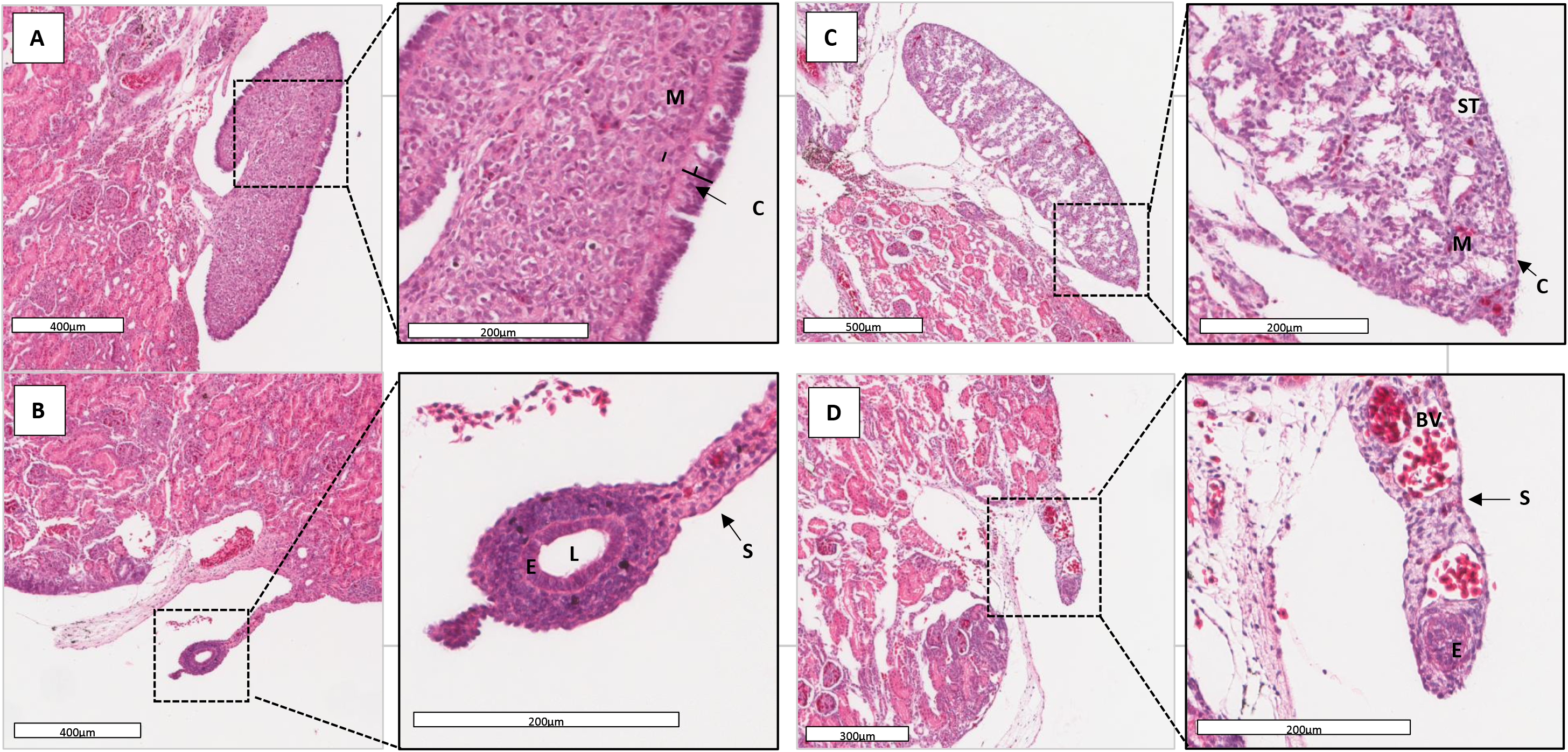
H & E stained cross-sections through (A) ovaries, (B) female Müllerian duct, (C) testis and (D) male Müllerian duct of a female and male green turtle hatchling. Images are x60 magnification and inserts x200 magnification. Female hatchlings had ovaries characterised by a thick cortex (C) and a medulla (M) consisting of dense mesenchymal tissue, and a long stalked (S) Müllerian duct consisting of a clear lumen (L) surrounded by a columnar epithelium (E). Male hatchlings had testes characterised by a thin cortex with differentiated seminiferous tubules (ST) within in the medulla, and a short stalked degenerating Müllerian duct often containing blood vessels (BV).

### Predicting hatchling sex ratio from incubation temperature

The constant temperature equivalents (CTE) during the SDP for treatments 3–10 were calculated using the temperature traces and these substituted in to the algorithm relating incubation temperature to sex-ratio (Fig. 1) to predict the hatchling sex-ratio of each treatment.

### Statistical analyses

Statistical analyses was performed using R version 4.0 (R Core Team 2020) and R Studio software (Rstudio Team 2020). For green turtle data, we used a generalised linear mixed-effects model (GLMM) using a binomial family (‘lme4’ package (Bates et al., 2015)) with two fixed factors: the incubation day then the temperature drop began (‘time’) (days 19, 23, 27 or 30 of incubation) and the temperature drop duration (‘days’) (3 or 7 days) and clutch as a random factor to test for differences of these factors on the proportion of males produced. Interactions that showed no significance were removed and the model was re-run to determine if drop period and duration independently influenced sex determination. We conducted a *post hoc* test analysis using Tukey’s Honest Significance Difference to test differences among the different drop times during the sex determining period. We used the same statistical procedures to analyse loggerhead turtle data, but because only one clutch of eggs was available, the clutch random factor was removed. We assumed significance at *P* < 0.05.

## RESULTS

### Hatching success

A total of 252 green turtle and 66 loggerhead turtle hatchlings were produced. The number of hatchlings per clutch for green turtles varied from 27-90. Hatching success across all treatments ranged from 49-69% for green turtle eggs and 44-89% for loggerheads, with an overall hatching success of 58% for green turtle eggs and 73% for loggerhead turtle eggs (Table 2).

**Table 2.** Hatching success and mean temperatures during the 26.50C temperature exposure for green turtle and loggerhead turtle eggs.

| Treatment | T1 | T2 | T3 | T4 | T5 | T6 | T7 | T8 | T9 | T10 |
| --- | --- | --- | --- | --- | --- | --- | --- | --- | --- | --- |
| Drop duration (days) | 0 | 0 | 3 | 7 | 3 | 7 | 3 | 7 | 3 | 7 |
| Drop time during incubation | 0 | 0 | 19 | 19 | 23 | 23 | 27 | 27 | 30 | 30 |
| Green turtle<br>(4 clutches) |  |  |  |  |  |  |  |  |  |  |
| Number of egg set | 46 | 45 | 44 | 44 | 43 | 42 | 42 | 42 | 42 | 42 |
| Number of eggs hatched | 27 | 22 | 25 | 24 | 21 | 27 | 24 | 27 | 26 | 29 |
| Hatching success (%) | 58.7 | 48.9 | 56.8 | 54.5 | 48.8 | 64.3 | 57.1 | 64.3 | 61.9 | 69.0 |
| Mean temperature during 26.5°C exposure (°C) | N/A | N/A | 26.9 | 26.7 | 27 | 26.8 | 27.1 | 26.9 | 27.3 | 27.0 |
| Loggerhead turtles<br>(1 clutch) |  |  |  |  |  |  |  |  |  |  |
| Number of egg set | 9 | 9 | 9 | 9 | 9 | 9 | 9 | 9 | 9 | 9 |
| Number of eggs hatched | 8 | 8 | 5 | 8 | 4 | 6 | 5 | 7 | 7 | 8 |
| Hatching success (%) | 88.9 | 88.9 | 55.6 | 88.9 | 44.4 | 66.7 | 55.6 | 77.8 | 77.8 | 88.9 |
| Mean temperature during 26.5°C exposure (°C) | N/A | N/A | N/A<sup>a</sup> | 26.8 | 27.1 | 26.8 | 26.7 | N/A<sup>a</sup> | 27.2 | 26.9 |

<sup>a</sup>Buttons in these containers stopped recording partway through incubation.

### Incubation temperature

Temperature traces from data loggers indicated similar temperatures to the constant temperature room set points (26.5°C and 30.0°C). The mean temperatures over the entire incubation period for treatments T1 and T2 in green turtle eggs were 27.1°C and 30.0°C, respectively. The temperature in treatment T1 of the loggerhead experiment increased above 26.5°C on day 39 of incubation due to failure of the refrigeration unit, so these eggs were moved into the 30.0°C room after day 39. The mean temperature for treatment T1 of loggerhead turtles was 26.5°C from day 1 of incubation until the refrigeration unit failed on day 39, and 28.3°C for the entire incubation period. The mean temperature for loggerhead turtle eggs incubated in treatment T2 was 30.0°C.

The mean incubation temperatures for the time that eggs were transferred to 26.5°C across all treatments ranged between 26.7–27.3°C and 26.7–27.2°C for green and loggerhead turtle eggs, respectively (Table 2). Temperature logger failure meant that temperature traces were not obtained from treatments T3 and T8 for loggerhead turtle eggs (Table 2).

For green turtle eggs, CTEs during the SDP for treatments T3-T10 remained above the upper TRT so no male hatchling production was predicted for these treatments (Table 3). In contrast, for loggerhead eggs, CTEs during the SDP for treatments T3-T10 remained within the TRT, so some male hatchling production was predicted (Table 3).

**Table 3.** Constant temperature equivalents (CTE) for the sex determining period (SDP), the proportion of males calculated using CTE and the incubation temperature – hatchling sex-ratio algorithm, and the empirically determined (by gonad histology) proportion of males for green and loggerhead turtle eggs incubated using the different incubation regimes. Green turtle temperature traces were used to predict the proportion of male hatchlings for the loggerhead treatments 3 and 8 where the temperature loggers failed.

| Species | Treatment | CTE |  |  |  |
| --- | --- | --- | --- | --- | --- |
|  |  | during<br>SDP (°C) | Predicted<br>males (%) | Empirical<br>males (%) | Sample<br>size |
| Green turtle | T1 | 26.5 | 100 | 100 | 27 |
|  | T2 | 30.1 | 0 | 9 | 22 |
|  | T3 | 29.7 | 0 | 12 | 25 |
|  | T4 | 29.2 | 0 | 8 | 24 |
|  | T5 | 29.6 | 0 | 5 | 21 |
|  | T6 | 29.2 | 0 | 30 | 27 |
|  | T7 | 29.6 | 0 | 29 | 24 |
|  | T8 | 29.2 | 0 | 48 | 27 |
|  | T9 | 29.7 | 0 | 8 | 26 |
|  | T10 | 29.0 | 0 | 34 | 29 |
| Loggerhead turtle | T1 | 28.6 | 44 | 75 | 8 |
|  | T2 | 30.5 | 10 | 13 | 8 |
|  | T3 | 30.1 | 14 | 60 | 5 |
|  | T4 | 29.5 | 24 | 88 | 8 |
|  | T5 | 30.3 | 12 | 25 | 4 |
|  | T6 | 29.4 | 26 | 33 | 6 |
|  | T7 | 30.0 | 15 | 20 | 5 |
|  | T8 | 29.4 | 26 | 29 | 7 |
|  | T9 | 30.2 | 13 | 71 | 7 |
|  | T10 | 30.1 | 14 | 75 | 8 |

### Effects of short-term drops on sex determination

Male hatchling production determined by histology of gonads was observed across all temperature drop commencement times (days 19, 23, 27, 30), for both durations (3 and 7 days) in both green and loggerhead turtle eggs (Table 3). In green turtle eggs, for treatments T1 (male control) and T2 (female control), the proportion of male hatchlings were 100% and 9%, respectively, while for loggerhead turtle eggs the proportion of males were 75% and 13% respectively (Table 3).

In green turtle experiments, the highest male hatchling production occurred when the temperature drop occurred on day 27 of incubation for seven days and the lowest when the temperature drop occurred on day 23 for three days (Fig. 4A). There was no interaction between the timing and duration of the temperature drop (P < 0.05). The proportion of males was always greater for the seven day drop treatments compared to the three day drop treatments (Fig. 4A, P < 0.001), and the proportion of males produced was greater when the temperature drop began on days 27 and 30 compared to day 19 (P = 0.001, and 0.023 respectively). Within green turtle clutches, clutch 4 had a much greater propensity (four times greater) to produce male hatchlings compared to the other three clutches sampled (Fig. 5).

**Figure 4.**
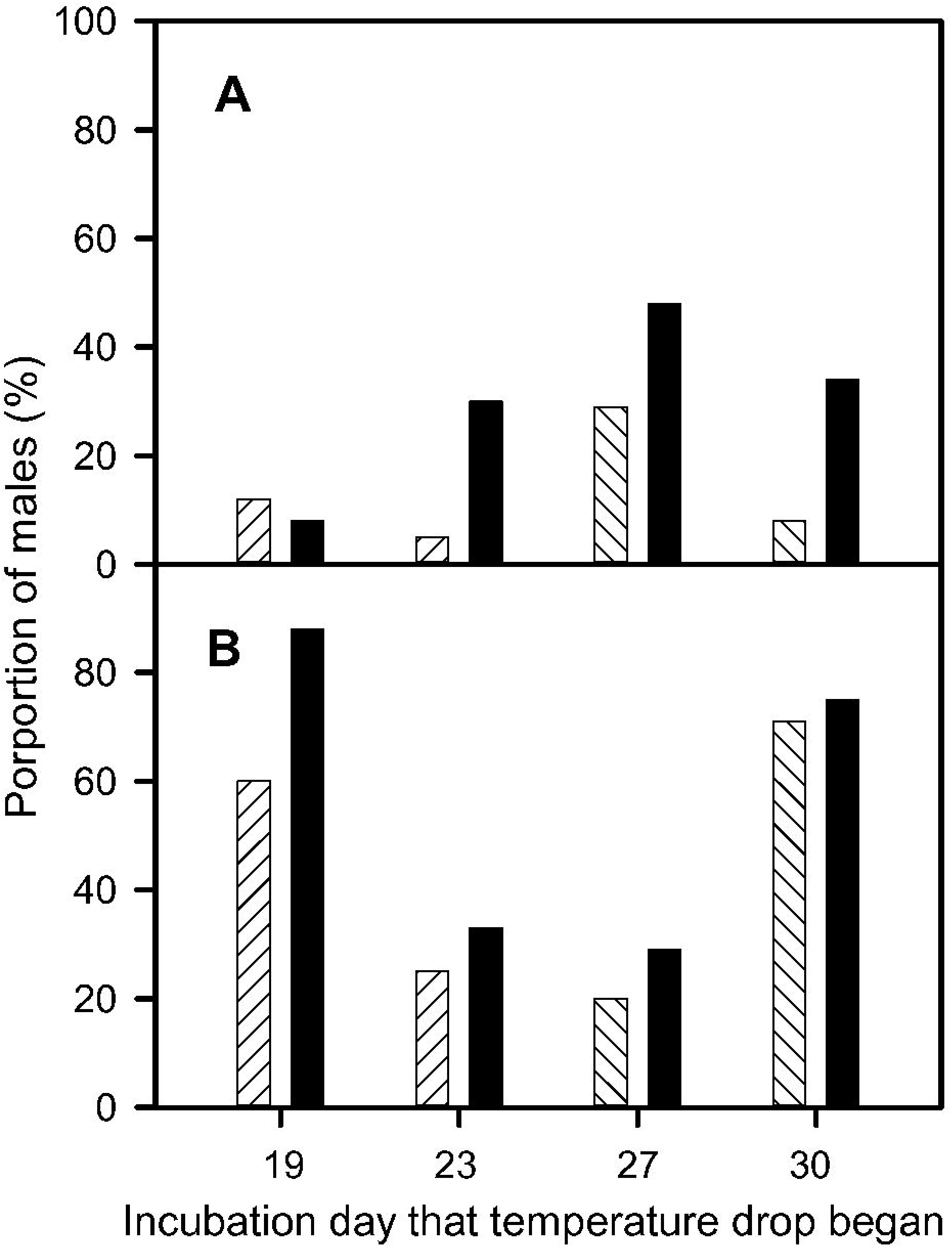
Proportion of male hatchlings as a function of both day of incubation that the temperature drop to 26.5°C began and the duration of the temperature drop. Striped bars represent a temperature drop duration of three days, solid bars represent a temperature drop duration of seven days. A. Green turtles, B. Loggerhead turtles.

**Figure 5.**
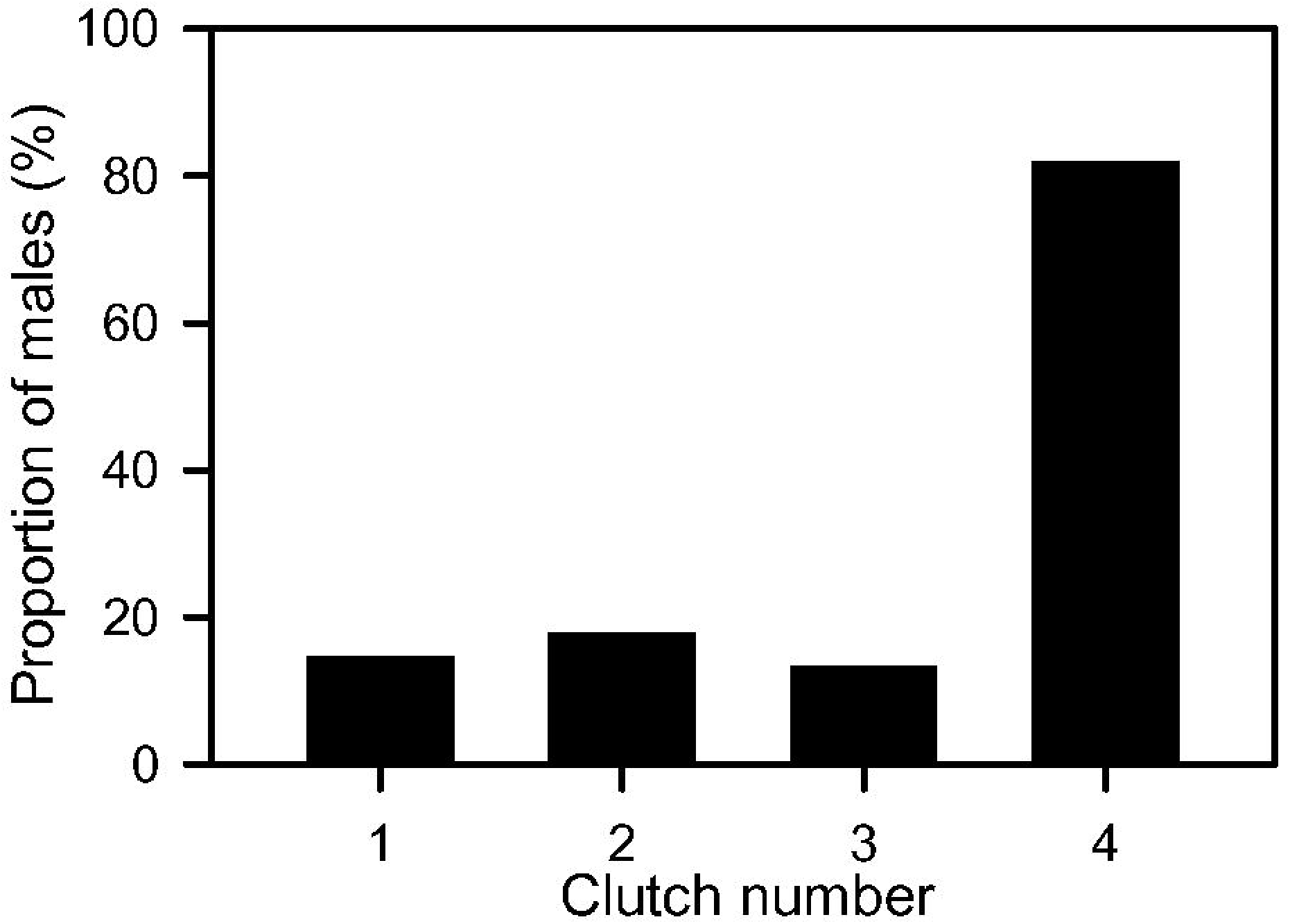
Proportion of male hatchlings produced pooled across treatments T3-T10 for the 4 different clutches of green turtle eggs examined.

In loggerhead turtle experiments, male hatchling production was highest when the temperature drop time occurred on day 19 of incubation for seven days and the lowest when the drop time occurred on day 27 for three days (Fig. 4B). There was no interaction between the timing and duration of the temperature drop (P < 0.05). The proportion of males was always greater for the seven day drop duration compared to the three day drop, but this difference was not statistically significant (Fig. 4B, P = 0.587), and more males were produced when the temperature drop began on days 19 and 30 compared to days 23 and 27 (Fig. 4B, P < 0.05).

## Discussion

Some male hatchling production occurred when eggs were exposed to short-term drops in temperature during the SDP in all of our temperature drop protocols, even when the drop duration was just three days. Previously, the shortest period of exposure to lower temperature to produce some male hatchlings was five days in a West Australian (WA) population of loggerhead turtles (Woolgar et al., 2013). This finding has important implications for sea turtle populations nesting at beaches where sand temperature is generally above the upper TRT so that only female hatchlings are thought to be produced. Numerous studies have predicted catastrophic feminisation of sea turtle populations within the next 50-100 years because of high nesting beach sand temperatures (e.g. Fuentes et al., 2009; Hawkes et al., 2009; Laloë et al., 2016; Jensen et al., 2018; Tanner et al,. 2019; Binhamer et al., 2019). Our findings indicate that at such beaches, heavy rainfall events that reduce nest temperature by 3°C for as short as three days can result in some male production. Previously, it was assumed that nest temperature needed to remain within the TRT throughout the SDP in order for some male production to occur, but clearly this is not the case. Prolonged rainy and cloudy weather has been associated with increased male sea turtle hatchling production (Reed 1980; Godfrey et al., 1996; Houghton et al., 2007; Lolavar and Wyneken, 2015; Wyneken and Lolavar, 2015; Lolavar and Wyneken 2017), but short-term drops in sand temperature have only recently been considered a possible mechanism to increase male hatchling production (Laloë et al., 2020; Staines et al., 2020). Our findings also make the possible use of irrigation to increase male hatchling production at managed sea turtle beaches easier because irrigation does not have to be applied throughout the SDP, once-only application maybe all that is required to produce some male hatchlings. Field trials are required to test the practicability of such interventions.

Our modelling using temperature traces to predict hatchling sex-ratio underestimated the proportion of males actually produced in all of our temperature drop treatments, so such modelling is a poor predictor of hatchling sex-ratios when eggs are exposed to short-term temperature drops during the SDP. Using temperature trace modelling to predict hatchling sex-ratio in natural sea turtle nests has been advocated as a method because it does not involve euthanising hatchlings. However, this type of modelling method is a poor predictor of actual sex-ratios in painted turtle (*Chrysemys picta*) (Carter et al., 2019) so it might also prove to be a poor predictor for natural sea turtle nests as well.

We found differences in the propensity to produce males between loggerhead and green turtle eggs, and between clutches in green turtles. In green turtle eggs, the seven-day drop in incubation temperature produced significantly more males than the three-day temperature drop, and the propensity to produce males was greatest during the middle of the SDP. One clutch also had a much greater propensity to produce males than the other three clutches. In the loggerheads turtle clutch, the duration of the temperature drop did not influence the propensity to produce males, and greater male production occurred at the beginning and the end of the SDP. A priori, we expected greater male production to be associated with longer temperature drop duration because embryos are exposed to male-producing temperatures for a longer period during SDP. Hence, the finding that propensity to produce loggerhead turtle embryos was similar for both temperature drop durations was unexpected, especially as longer temperature drop duration was associated with greater male production in a West Australian (WA) nesting population of loggerhead turtles (Woolgar et al., 2013). The small sample size probably explains our result, because the proportion of males produced was slightly greater for the longer duration across all times within the SDP, but this difference was not great enough to reach statistical significance. There are also differences between the East Australian (EA) and WA loggerhead turtle nesting populations with respect to the influence of timing within the SDP for the propensity to produce males, and the pivotal temperate and TRT. In contrast to our findings, in the WA loggerhead turtle population the propensity to produce males was similar throughout the SDP, and the pivotal temperature is higher (29.0°C WA Vs 28.4°C EA) and the TRT much narrower (28.7 - 29.3°C WA Vs 25.0 – 32°C EA) (Woolgar et al., 2013). The mechanism underlying differences in timing within the SDP in the propensity to produce males between green and loggerhead turtle embryos is unknown, but has important implications if once-off artificial irrigation of nests is to be used as a management tool to increase male hatchling production. In the populations we studied, such an intervention should be targeted for the middle of the SDP in green turtle nests, and towards the beginning or end of the SDP in loggerhead turtle nests. Inter-clutch differences in the propensity to produce male hatchlings when developing embryos experience the same incubation regime (i.e. differences in pivotal temperature and TRT), have been reported previously in sea turtles (Limpus et al., 1985; Mrosovsky, 1988; Wibbles, 2003; Lockley et al., 2020) and freshwater turtles (Bobyn and Brooks, 1994; Bowden et al., 2001; Dodd et al., 2006; Carter et al., 2019). These inter-population and inter-clutch differences in the relationship between incubation temperature and hatchling sex-ratios may be caused by genetic factors that influence the developmental rate of the gonads and/or by maternal egg-specific factors such as yolk steroid hormone concentrations (Bowden et al. 2001, Dodd et al. 2006, Díaz-Hernández et al. 2015, Lockley et al. 2020). For example, in loggerhead turtles, low concentrations of oestradiol (E_2_) and testosterone (T) within the yolk at laying were associated with greater male production, and higher concentration of either E_2_ or T were associated with greater female production (Lockley et al. 2020). The inter-clutch variation in the propensity to produce males combined with short-term temperature drop-induced male production may become an important avenue for male hatchling production at sea turtle nesting beaches where climate warming of sand is causing a general feminisation of hatchlings.

Our loggerhead male producing control treatment produced a relatively large number of female hatchlings. This result can be explained by two phenomena. First, the EA nesting population of loggerhead turtles has a particular wide TRT, with some female production expected at temperatures as low as 25 °C (Limpus et al., 1985). Second, the failure of the refrigeration unit on day 39 of incubation in this treatment meant that embryos were exposed to female producing temperatures after day 39. The SDP for loggerhead turtle embryos incubated at 26.5°C was expected to occur between day 22 and day 46 of incubation, so exposure to female-producing temperatures after day 39 may have caused an increase in female hatchling production. Some male hatchlings were also produced at the constant 30.0°C temperature, and once again, this can be explained by the wide TRT for this population where some female production is expect at temperatures up to 32°C (Limpus et al., 1985).

While further investigation is necessary to pinpoint precisely when male hatchling production occurs during the SDP, our study revealed that male hatchlings can be produced when exposed to temperature drops for as little as 3 days during the SDP. This knowledge indicates that some male hatchling production is likely to occur at otherwise all female producing nesting beaches if occasional heavy rainfall events occur during the nesting season. It also has the potential to allow sea turtle nesting beach managers to increase male hatchling production through the use of once-off artificial irrigation to cool nest temperatures for a short period of time. The use of one-off irrigation will save on water resources, compared to the previously assumption that multiple irrigation events were required in order to maintain nest temperature below all female producing temperatures throughout the SDP.

## Acknowledgements

We thank staff and volunteers, particularly Lizzy Turner at Mon Repos Conservation and the staff at Heron island research station for their assistance with collecting eggs. We thank Caitlin Smith (WWF) and Melissa Staines (UQ) for participating in the blind identifying hatchling sex from gonadal sections. We thank Simone Blomberg for advice and assistance with statistical analyses.

## Author contributions

Conceptualization: E.P., D.T.B., C.J.L.; Methodology: E.P., D.T.B., C.J.L.; Formal analysis: E.P.; Investigation: E.P.; Resources: D.T.B., C.J.L.; Data curation: E.P.; Writing - original draft: E.P., D.T.B; Writing - review & editing: E.P., D.T.B., Visualization: E. P. D.T.B.; Supervision: D.T.B., C.J.L; Project administration: D.T.B.; Funding acquisition: D.T.B.

## References

Ackerman, R. A. (1997). The nest environment and the embryonic development of sea turtles. In The Biology of Sea Turtles (ed. P. L. Lutz and J. A. Musick), pp. 83–106. Boca Raton, Florida: CRC Press.

Ackerman, R. A., and Lott, D.B. (2004). Thermal, hydric and respiratory climate of nests. In Reptilian Incubation. Environment, Evolution and Behaviour (ed. D. C. Deeming), pp. 15–43. Nottingham: Nottingham University Press.

Bancroft, J. D., and Layton, C. (2012). The hematoxylins and eosin. In Bancroft’s Theory and Practice of Histological Techniques (ed. J. D. Bancroft), pp. 173–186. London: Churchill Livingstone.

Bates, D., Mächler, M., Bolker, B., and Walker, S. (2015). Fitting linear mixed-effects models using lme4. J. Stat. Soft. 67, 1–48.

Binhammer, M. R., Beange, M., and Arauz, R. (2019). Sand temperature, sex ratios, and nest success in olive ridley sea turtles. Mar. Turt. News. 159, 5–9.

Bobyn, M. L., and Brooks, R.J. (1994). Interclutch and interpopulation variation in the effects of incubation conditions on sex, survival and growth of hatchling turtles (*Chelydra serpentina*). J. Zool. 233, 233–257.

Booth, D. T. (2006). Influence of incubation temperature on hatchling phenotype in reptiles. Physiol. Biochem. Zool. 79, 274–281.

Booth, D. T. (2017). Influence of incubation temperature on sea turtle hatchling quality. Integr. Zool. 12, 352–360.

Booth, D. T., and Astill, K. (2001a). Incubation temperature, energy expenditure and hatchling size in the green turtle (*Chelonia mydas*), a species with temperature-sensitive sex determination. Aust. J. Zool.: 49, 389–396.

Booth, D. T., and Astill, K. (2001b). Temperature variation within and between nests of the green sea turtle, *Chelonia mydas* (Chelonia: Cheloniidae) on Heron Island, Great Barrier Reef. Aust. J. Zool. 49, 71–84.

Booth, D. T., and Freeman, C. (2006). Sand and nest temperatures and an estimate of hatchling sex ratio from the Heron Island green turtle (*Chelonia mydas*) rookery, Southern Great Barrier Reef. Coral Reefs 25, 629–633.

Booth, D.T., Burgess, E., McCosker, J., and Lanyon, J.M. (2004). The influence of incubation temperature on post-hatching fitness characteristics of turtles. Inter. Cong. Ser. 1275, 226–233.

Booth, D.T., Dunstan, A., Bell, I., Reina, R., Tedeschi, J. (2020). Low male production at the world’s largest green turtle rookery. Mar. Ecol. Prog. Ser. 653, 181–190. https://doi.org/10.3354/meps13500.

Bowden, R. M., Ewert, M. A., Lipar, J. L., and Nelson, C. E. (2001). Concentrations of steroid hormones in layers and biopsies of chelonian egg yolks. Gen. Comp. Endo. 121, 95–103.

Broderick, A., Godley, B., Reece, S., and Downie, J. (2000). Incubation periods and sex ratios of green turtles: highly female biased hatchling production in the eastern Mediterranean. Mar. Ecol. Prog. Ser. 202, 273–281.

Broderick, A. C., Godley B. J., and Hays, G.C. (2001) Metabolic heating and the prediction of sex ratios for green turtles (*Chelonia mydas*). Physiol. Biochem. Zool. 74, 161–170.

Burgess, E. A., Booth, D. T., and Lanyon, J. M. (2006). Swimming performance of hatchling green turtles is affected by incubation temperature. Coral Reefs 25, 341–349.

Bustard, H. R., and Greenham, P. (1968). Physical and chemical factors affecting hatching in green sea turtle *Chelonia mydas* (L). Ecology 49, 269–276.

Carter, A. L., Bodensteiner, B. L., Iverson, J. B., Milne-Zelman, C. L., Mitchell, T. S., Refsnider, J. M., Warner, D. A., and Janzen, F. J. (2019). Breadth of the thermal response captures individual and geographic variation in temperature-dependent sex determination. Funct. Ecol. 33, 1928–1939.

Ceriani, S. A., and Wyneken, J. (2008). Comparative morphology and sex identification of the reproductive system in formalin-preserved sea turtle specimens. Zoology 111, 179–187.

Chu, C., Booth, D. T., and Limpus, C. J. (2008). Estimating the sex ratio of loggerhead turtle hatchlings at Mon Repos rookery (Australia) from nest temperatures. Aust. J. Zool. 56, 57–64.

Díaz-Hernández, V., Marmolejo-Valencia, A., and Merchant-Larios, H. (2015). Exogenous estradiol alters gonadal growth and timing of temperature sex determination in gonads of sea turtle. Develop. Biol. 408, 79–89.

Dodd, K. L., Murdock, C., and Wibbels, T. (2006). Interclutch variation in sex ratios produced at pivotal temperature in the red-eared slider, a turtle with temperature-dependent sex determination. J. Herpetol. 40, 544–550.

Esteban, N., Laloë, J. O., Kiggen, F. S. P. L. S., Ubels, M., Becking, L.E., Meesters, E. H., Berkel, J., Hays, G. C., and Christianen, M. J. A. (2018). Optimism for mitigation of climate warming impacts for sea turtles through nest shading and relocation. Sci. Reports 8, 17625.

Ewert, M. A., Jackson, D. R., and Nelson, C. E. (1994). Patterns of temperature‐dependent sex determination in turtles. J. Exper. Zool. 270, 3–15.

Fisher, L.R., Godfrey, M.H., Owens, D.W. (2014). Incubation temperature effects on hatchling performance in the loggerhead sea turtle (*Caretta caretta*). PloS ONE 9, e114880.

Fuentes, M.M.P.B., Maynard, J., Guinea, M., Bell, I., Werdell, P., and Hamann, M. (2009). Proxy indicators of sand temperature help project impacts of global warming on sea turtles in northern Australia. Endang. Species Res. 9, 33–40.

Fuentes, M.M.P.B., Fish, M.R., Maynard, J.A. (2012). Management strategies to mitigate the impacts of climate change on sea turtle’s terrestrial reproductive phase. Mit. Adapt. Strat. Glob. Chan. 17, 51–63.

Georges, A., Limpus, C., and Stoutjesdijk, R. (1994). Hatchling sex in the marine turtle *Caretta caretta* is determined by proportion of development at a temperature, not daily duration of exposure. J. Exper. Zool. 270, 432–444.

Girondot, M. (1999). Statistical description of temperature-dependent sex determination using maximum likelihood. Evol. Ecol. Res. 1, 479–486.

Godfrey, M. H., and Mrosovsky, N. (1999). Estimating hatchling sex ratios. In Research and Management Techniques for the Conservation of Sea Turtles (ed. K. L. Eckert, K. A. Bjorndal, F. A. Abreu-Grobois and M. Donnelly), pp. 136–138. Gland, Switzerland: Marine Turtle Specialist Group, IUCN/SSC.

Godfrey, M. H., and Mrosovsky, N. (2006). Pivotal temperature for green sea turtles, *Chelonia mydas*, nesting in Suriname. Herpetol. J. 16, 55–61.

Godfrey, M. H., Mrosovsky, N., and Barreto, R. (1996). Estimating past and present sex ratios of sea turtles in Suriname. Can. J. Zool. 74, 267–277.

Godley, B., Broderick, A., Downie, J., Glen, F., Houghton, J., Kirkwood, I., Reece, S., and Hays, G. (2001). Thermal conditions in nests of loggerhead turtles: further evidence suggesting female skewed sex ratios of hatchling production in the Mediterranean. J. Exper. Mar. Biol. Eco. 263, 45–63.

Hawkes, L. A., Broderick, A. C., Godfrey, M. H., and Godley, B. J. (2009). Climate change and marine turtles. Endang. Species Res. 7, 137–154.

Hays, G., Ashworth, J., Barnsley, M., Broderick, A., Emery, D., Godley, B., Henwood, A., and Jones, E. (2001). The importance of sand albedo for the thermal conditions on sea turtle nesting beaches. Oikos 93, 87–94.

Hill, J. E., Paladino, F. V., Spotila, J. R., and Tomillo, P. S. (2015). Shading and watering as a tool to mitigate the impacts of climate change in sea turtle nests. PloS One 10:e0129528.

Houghton, J. D. R., Myers, A. E., Lloyd, C., King, R. S., Isaacs, C., and Hays, G. C. (2007). Protracted rainfall decreases temperature within leatherback turtle (*Dermochelys coriacea*) clutches in Grenada, West Indies: Ecological implications for a species displaying temperature dependent sex determination. J. Exper. Mar. Biol. Ecol. 345, 71–77.

Jensen, M. P., Allen, C. D., Eguchi, T., Bell, I. P., LaCasella, E. L., Hilton, W. A., Hof, C. A., and Dutton, P. H. (2018). Environmental warming and feminization of one of the largest sea turtle populations in the world. Current Biol. 28, 154–159.

Jourdan, J., and Fuentes, M. (2015). Effectiveness of strategies at reducing sand temperature to mitigate potential impacts from changes in environmental temperature on sea turtle reproductive output. Miti. Adapt. Strat. Glob. Chang. 20, 121–133.

Katselidis, K.A., Schofield, G., Stamou, G., Dimopoulos, P., Pantis, J. D. (2012). Females first? Past, present and future variability in offspring sex ratio at a temperate sea turtle breeding area. Anim. Conser. 15, 508–518.

King, R., Cheng, W. H., Tseng, C. T., Chen, H., and Cheng, I. J. (2013). Estimating the sex ratio of green sea turtles (*Chelonia mydas*) in Taiwan by the nest temperature and histological methods. J. Exper. Mar. Biol. Ecol. 445, 140–147.

Laloë, J. O., Esteban, N. Berkel, J., and Hays, G. C. (2016). Sand temperatures for nesting sea turtles in the Caribbean: implications for hatchling sex ratios in the face of climate change. J. Exper. Mar. Biol. Ecol. 474, 92–99.

Laloë, J.-O., Tedeschi, J.N., Booth, D.T., Bell, I., Dunstan, A. Reina, R.D., Hays, G.C. (2020). Extreme rainfall events and cooling of sea turtle clutches: Implications in the face of climate warming. Ecol. Evol. Doi: 10.1002/ece3.7076

Limpus, C. J., Reed, P. C., and Miller, J. D. (1985). Temperature dependent sex determination in Queensland sea turtles: intraspecific variation in *Caretta caretta*. In Biology of Australian Frogs and Reptiles (ed. G. Grigg, R. Shine and H. Ehmann.), pp. 343–351. Sydney: Surrey Beatty and Sons.

Lockley, E. C., Reischig, T., and Eizaguirre, C. (2020). Maternally derived sex steroid hormones impact sex ratios of loggerhead sea turtles. bioRxiv. doi: https://doi.org/10.1101/2020.01.10.901520

Lolavar, A., and Wyneken, J. (2015). Effect of rainfall on loggerhead turtle nest temperatures, sand temperatures and hatchling sex. Endang. Spec. Res. 28, 235–247.

Lolavar, A., and Wyneken, J. (2017). Experimental assessment of the effects of moisture on loggerhead sea turtle hatchling sex ratios. Zoology 123, 64–70.

Miller, J. D. (1982). Development of marine turtles. PhD thesis, University of New England, Armidale, NSW, Australia.

Miller, J. D. (1997). Reproduction in sea turtles. In The Biology of Sea Turtles (ed. P. L. Lutz and J. A. Musick), pp. 51–81. Boca Raton, Florida:. CRC Press.

Miller, J., and Limpus, C. (1981). Incubation period and sexual differentiation in the green turtle *Chelonia mydas*. In Proceedings of the Melbourne Herpetological Symposium (ed. C. B. Banks and A. A. Martin), pp. 66–73. Melbourne: The Zoological Board of Victoria.

Mrosovsky, N. (1988). Pivotal temperatures for loggerhead turtles (*Caretta caretta*) from northern and southern nesting beaches. Can. J. Zool. 66, 661–669.

Mrosovsky, N. (2006). Distorting gene pools by conservation: assessing the case of doomed turtle eggs. Environ. Manage. 38, 703–703.

Mrosovsky, N., and Provancha, J. (1992). Sex ratio of hatchling loggerhead sea turtles: data and estimates from a 5-year study. Can. J. Zool. 70, 530–538.

Ozdemir, A., Ilgaz, C., Durmus, S.H., Guclu, O. (2011). The effect of the predicted air temperature change on incubation temperature, incubation duration, sex ratio and hatching success of loggerhead turtles. Anim. Biol. 61, 369–81.

Parmenter, C. (1980). Incubation of the eggs of the green sea turtle, *Chelonia mydas*, in Torres Strait, Australia: the effect of movement on hatchability. Wildl. Res. 7, 487–491.

Patino‐Martinez, J., Marco, A., Quiñones, L., and Hawkes, L. (2012). A potential tool to mitigate the impacts of climate change to the Caribbean leatherback sea turtle. Global Chang. Biol. 18, 01–411.

R Core Team. (2020). R: a language and environment for statistical computing. R Foundation for Statistical Computing, Vienna, Austria. URL: https://www.R-project.org/.

RStudio Team. (2020). RStudio: integrated development environment for R. RStudio, PBC, Boston, Massachusetts, U.S.A. URL: http://www.rstudio.com/.

Reboul, I., Booth D., and Rulsi U. (2021). Artificial and natural shade: Implications for green turtle (*Chelonia mydas*) rookery management. Ocean Coast. Manag. 204, https://doi.org/10.1016/j.ocecoaman.2021.105521

Reed, P. C. (1980). The sex ratios of hatchling loggerhead turtles. Honours Thesis. James Cook University, Queensland, Australia.

Rivas, M. L., Esteban, N., and Marco, A. (2019). Potential male leatherback hatchlings exhibit higher fitness which might balance sea turtle sex ratios in the face of climate change. Climatic Change 156, 1–14.

Rivas, M. L., Spínola, M., Arrieta, H., and Faife‐Cabrera, M. (2018). Effect of extreme climatic events resulting in prolonged precipitation on the reproductive output of sea turtles. Anim. Cons. 21, 387–395.

Saba, V.S., Stock, C.A., Spotila, J.R., Paladino, F. V., Tomillo, P. S. (2012). Projected response of an endangered marine turtle population to climate change. Nat. Clim. Chang. 2, 814–20.

Sari, F., and Kaska, Y. (2016). Histochemical and immunohistochemical studies of the gonads and paramesonephric ducts of male and female hatchlings of loggerhead sea turtles (*Caretta caretta*). Biotech. Histochem 91, 428–437.

Sim, E., Booth, D., and Limpus, C. (2015). Incubation temperature, morphology and performance in loggerhead (*Caretta caretta*) turtle hatchlings from Mon Repos, Queensland, Australia. Biol. Open 4, 685–692.

Spotila, J. R. (2004). Sea Turtles: A Complete Guide to Their Biology, Behavior, and Conservation. John Hopkins University Press, Baltimore, Maryland, U.S.A.

Spotila, J. R., Standora, E. A., Morreale, S. J., and Ruiz, G. J. (1987). Temperature-dependent sex determination in the green turtle (*Chelonia mydas*): effects on the sex ratio on a natural nesting beach. Herpetologica 1, 74–81.

Staines, M. N., Booth, D. T., and Limpus, C. J. (2019). Microclimatic effects on the incubation success, hatchling morphology and locomotor performance of marine turtles. Acta Oecol. 97, 49–56.

Staines, M. N., Booth, D. T., Hof, C. A. M., and Hays, G. C. (2020). Impact of heavy rainfall events and shading on temperature of sea turtle nests. Mar. Biol. 167, 190.

Standora, E. A., and Spotila, J. R. (1985). Temperature dependent sex determination in sea turtles. Copeia 1985, 711–722.

Tanner, C. E., Marco A., Martins, S., Abella-Perez, E., and Hawkes, L. A. (2019). Highly feminised sex-ratio estimations for the world’s third-largest nesting aggregation of loggerhead sea turtles. Mar. Ecol. Prog. Ser. 621, 209–219.

Tezak, B., Sifuentes-Romero, S., Milton, S., and Wyneken, J. (2020). Identifying sex of neonate turtles with temperature-dependent sex determination via small blood samples. Scientific Rep. 10, 1–8.

Weber, S.B., Broderick, A.C., Groothuis, T.G.G., Ellick, J., Godley, B. J., Blount, J. D. (2012). Fine-scale thermal adaptation in a green turtle nesting population. Proc. Roy. Soc 279, 1077–1084.

Wibbels, T. (2003). Critical approaches to sex determination in sea turtles. In The Biology of Sea Turtles (Volume II). (ed. P. L. Lutz, J. A. Musick and J. Wyneken), pp. 103–134). Boca Raton, Florida: CRC Press.

Wood, A., Booth, D. T., and Limpus, C. J. (2014). Sun exposure, nest temperature and loggerhead turtle hatchlings: implications for beach shading management strategies at sea turtle rookeries. J. Exper. Mar. Biol. Ecol. 451, 105–114.

Woolgar, L., Trocini, S., and Mitchell, N. (2013). Key parameters describing temperature-dependent sex determination in the southernmost population of loggerhead sea turtles. J. Exper. Mar. Biol. Ecol. 449, 77–84.

Wyneken, J., and Lolavar, A. (2015). Loggerhead sea turtle environmental sex determination: implications of moisture and temperature for climate change based predictions for species survival. J. Exper. Zool. B 324, 295–314.

Yntema, C., and Mrosovsky, N. (1980). Sexual differentiation in hatchling loggerheads (*Caretta caretta*) incubated at different controlled temperatures. Herpetologica 1, 33–36.

Yntema, C., and Mrosovsky, N. (1982). Critical periods and pivotal temperatures for sexual differentiation in loggerhead sea turtles. Can. J. Zool. 60, 1012–1016.

